# Challenges for targeting SARS-CoV-2 proteases as a therapeutic strategy for COVID-19

**DOI:** 10.1101/2020.11.21.392753

**Authors:** Kas Steuten, Heeyoung Kim, John C. Widen, Brett M. Babin, Ouma Onguka, Scott Lovell, Oguz Bolgi, Berati Cerikan, Mirko Cortese, Ryan K. Muir, John M. Bennett, Ruth Geiss-Friedlander, Christoph Peters, Ralf Bartenschlager, Matthew Bogyo

**Affiliations:** Department of Pathology, Stanford University School of Medicine, 300 Pasteur Drive, Stanford, California 94305, United States; Department of Infectious Diseases, Molecular Virology, University Hospital Heidelberg, Heidelberg, Germany; Institute of Molecular Medicine and Cell Research, University of Freiburg, Freiburg, Germany; Division Virus-Associated Carcinogenesis, German Cancer Research Center (DKFZ), Heidelberg, Germany; German Center for Infection Research (DZIF), Heidelberg partner site, Heidelberg, Germany

**Keywords:** SARS-CoV-2, Main Protease, Papain-Like Protease, Cathepsin Cross-Reactivity, 11a, GC376, Rupintrivir, COVID-19 Treatment

## Abstract

Two proteases produced by the SARS-CoV-2 virus, M^pro^ and PL^pro^, are essential for viral replication and have become the focus of drug development programs for treatment of COVID-19. We screened a highly focused library of compounds containing covalent warheads designed to target cysteine proteases to identify new lead scaffolds for both M^pro^ and PL^pro^ proteases. These efforts identified a small number of hits for the M^pro^ protease and no viable hits for the PL^pro^ protease. Of the M^pro^ hits identified as inhibitors of the purified recombinant protease, only two compounds inhibited viral infectivity in cellular infection assays. However, we observed a substantial drop in antiviral potency upon expression of TMPRSS2, a transmembrane serine protease that acts in an alternative viral entry pathway to the lysosomal cathepsins. This loss of potency is explained by the fact that our lead M^pro^ inhibitors are also potent inhibitors of host cell cysteine cathepsins. To determine if this is a general property of M^pro^ inhibitors, we evaluated several recently reported compounds and found that they are also effective inhibitors of purified human cathepsin L and B and showed similar loss in activity in cells expressing TMPRSS2. Our results highlight the challenges of targeting M^pro^ and PL^pro^ proteases and demonstrate the need to carefully assess selectivity of SARS-CoV-2 protease inhibitors to prevent clinical advancement of compounds that function through inhibition of a redundant viral entry pathway.

## INTRODUCTION

The emergence of the novel coronavirus, SARS-CoV-2 in late December 2019^1^ created a global pandemic, which has prompted unprecedented efforts to combat the virus using diverse vaccine and therapy strategies. One of the more promising therapeutic approaches involves repurposing existing drugs that can be rapidly advanced into clinical studies. Other strategies build on existing knowledge and lead molecules that were developed in response to earlier coronavirus outbreaks^2^. Two promising targets that emerged from the SARS-CoV-1 outbreak in 2003 were the essential main protease (M^pro^) and papain-like protease (PL^pro^)^3^. These two cysteine proteases are encoded in the viral polyprotein as non-structural protein (Nsp) 3 and Nsp5. They are responsible for cleavage of the viral polyprotein into several structural and non-structural proteins prior to formation of the replication organelle that is established in close proximity to virus assembly sites^3^. Therefore, inhibition of one or both of these enzymes effectively blocks viral RNA replication and thus virus transmission.

Several covalent inhibitors containing various electrophilic warheads including α-ketoamides, aldehydes and α,β-unsaturated ketones have been developed as inhibitors of M^pro^, with α-hydroxy ketone PF-07304814 recently entering human clinical trials (ClinicalTrials.gov, NCT04535167)^4^. The development of covalent small molecule inhibitors of PL^pro^ has been more challenging, perhaps due to a dominant non-proteolytic function and preference for relatively large ubiquitin-like protein substrates^5,6^. This premise is further supported by the fact that, while some small peptide-based inhibitors have been reported^7^, the most successful inhibitors target exosites involved in ubiquitin recognition^6,8,9^. While both M^pro^ and PL^pro^ are considered to be promising therapeutic targets, several properties of these proteases, combined with the past history of efforts to develop protease inhibitors for other RNA viruses such as hepatitis C virus (HCV)^10^ portend multiple challenges for drug discovery efforts. Like other proteases from RNA viruses, the M^pro^ protease liberates itself from a large polyprotein through N-terminal autocleavage before the mature, active dimer can be formed^11–13^. This initial event is difficult to inhibit due to the favorability of the intramolecular reaction. After maturation, the dimeric protease is likely localized to defined regions inside the cytosol or at membrane surfaces in proximity to its viral protein substrates resulting in relatively high local substrate concentrations. In addition, a number of viral proteases have been found to undergo product inhibition where they retain their cleaved substrates within the active site, thus requiring displacement for effective inhibitor binding^14,15^. Additionally, inhibition of M^pro^ prior to formation of its semi-active monomer is likely impossible due to the fact that this early stage intermediate lacks a properly formed active site^11,12,16^. Thus, inhibitors must be highly bioavailable and cell permeant such that they can reach local concentrations that are sufficient to compete with native substrates and inhibit the viral protease early in the infection cycle.

Another significant challenge for targeting M^pro^ and PL^pro^ is the potential for any lead molecule to target host proteases with similar substrate preferences. This is compounded by the diverse set of cellular systems used to evaluate lead molecules, which express different levels and types of proteases. There also remains controversy about which cell type best represents primary sites of infection *in vivo*^17,18^. In particular, priming of the receptor binding domain (RBD) of the S-glycoprotein of SARS-CoV-2 by host proteases is required after binding to the angiotensin converting enzyme-2 (ACE2) entry receptor^19^. This process can be mediated by multiple proteases including cysteine cathepsins B and L or the transmembrane protease serine 2 (TMPRSS2)^20,21^. While high expression levels of both cathepsins and TMPRSS2 have been confirmed in lung tissue^22^, cell lines commonly used for viral infection assays have varying expression levels of both protease classes which can have a dramatic impact on the mechanism used by the virus for entry^18^. The redundancy of these pathways not only poses a challenge for antiviral drugs that are targeted towards host factors such as cathepsins or TMPRSS2 (K11777, E64d or camostat^23–25^), but also for drugs that display off-target activity towards these enzymes.

In this work, we screened a highly focused library of ~650 cysteine reactive molecules against PL^pro^ and M^pro^ using a fluorogenic substrate assay to identify novel lead molecules as potential antiviral agents. From this screen, we identified six inhibitors containing various electrophiles, which demonstrated time-dependent inhibition of recombinant M^pro^. Notably, we did not identify any viable hits for PL^pro^. Two of the six lead M^pro^ inhibitors were active in cellular infectivity assays using A549 epithelial lung cells, but their potency decreased significantly upon expression of TMPRSS2 as was the case for established cysteine cathepsin inhibitors (E64d and K11777) and multiple previously reported M^pro^ inhibitors. This loss of potency could be best explained by the fact that TMPRSS2 expression provides an alternate entry pathway for the virus and therefore any lost antiviral activity was likely mediated by cathepsin inhibition. Indeed, we confirm cathepsin cross-reactivity of our newly discovered M^pro^ inhibitors as well as for several of the reported M^pro^ inhibitors. These results highlight the challenges for selection of M^pro^ inhibitors based on antiviral activity without complete understanding of their target selectivity as it can result in advancement of compounds based on disruption of redundant entry pathways rather than on direct antiviral effects.

## RESULTS AND DISCUSSION

To identify potential inhibitors of M^pro^ and PL^pro^, we developed fluorogenic substrate assays that allowed us to screen a focused library of cysteine reactive molecules. We based the design of internally quenched-fluorescent M^pro^ substrates on recent specificity profiling of the P1-4 residues using non-natural amino acids with a C-terminal 7-amino-4-carbamoylmethylcoumarin (ACC) reporter^26^. However, because the reported structures have relatively low turnover rates, we decided to make extended versions of these substrates that combine the optimal P1-4 residues with the native cleavage consensus of the P1’-P3’ residues (i.e., residues C-terminal of the scissile bond)^26,27^. This required synthesis of substrates using a quencher/fluorophore pair rather than an ACC reporter (Fig. 1A; Fig S1). A dramatic increase was observed in the catalytic rate of substrate conversion by M^pro^ as we incorporated more prime site residues into the substrate sequence (Fig 1B-C). This result explains the reason for the overall low kinetic rate constants for reported ACC substrates which lack any prime side residues. As a substrate for the PL^pro^ protease, we synthesized the reported ACC peptide derived from the ubiquitin consensus sequence LRGG (N-terminal acetylated substrate referred to as: Ac-LRGG-ACC)^7^. For activity assays we used recombinant M^pro^ and PL^pro^ that were cloned for expression in *E. coli* and subsequently purified (Fig S2A-B). We then optimized enzyme and substrate concentrations such that the Z-factors for each assay were consistently above 0.5. We found that substrate turnover by PL^pro^ required the presence of reducing agent DTT whereas it could be omitted in the M^pro^ assay (Fig S3).

**Figure 1.**
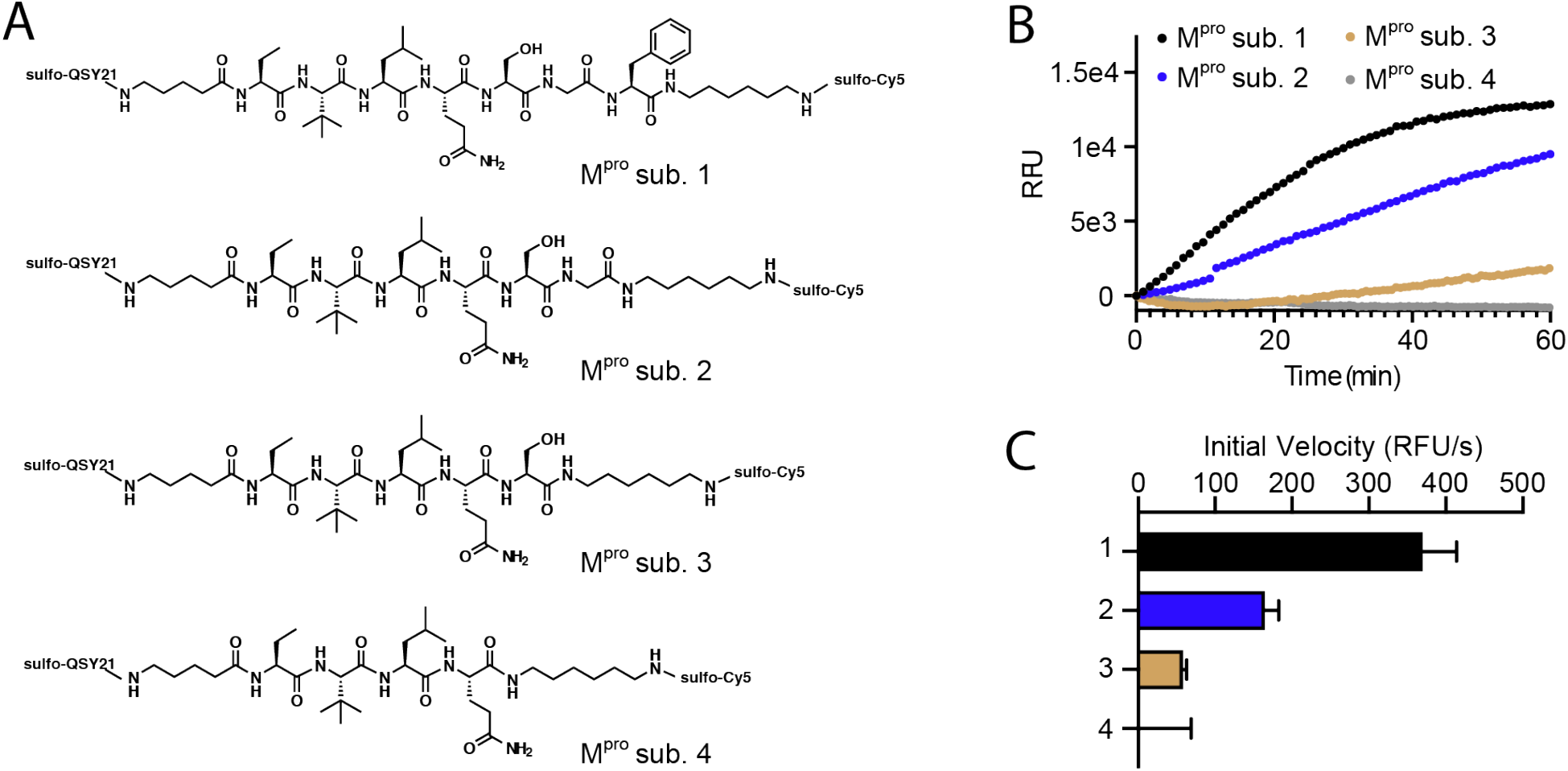
Design of quenched-fluorescent M^pro^ substrates for the inhibitor screening assay. A) Chemical structures of internally quenched M^pro^ substrates. B) Progress curves and C) initial velocities of M^pro^ substrates. 10 μM substrate was added to 100 nM M^pro^ immediately prior to fluorescence readout.

After having established optimal assay conditions, we screened a library of approximately 650 compounds designed to inhibit cysteine proteases^28,29^. Because this set of compounds contains a diverse but highly focused set of cysteine-reactive molecules, we have found that it produces viable lead scaffolds for virtually all the cysteine protease targets that we have screened. The library contains molecules with electrophiles including aza-peptide epoxyketones, aza-peptide vinylketones, epoxides, halomethylketones, acyloxymethylketones and sulfones. We screened the library by measuring residual enzymatic activity after 30 min incubation of M^pro^ substrate 2 and Ac-LRGG-ACC for PL^pro^. We set a threshold of maximum 10% residual M^pro^ activity and identified 27 hits. In subsequent time-dependent inhibition assays, the hits were further narrowed down to six validated reproducible covalent M^pro^ inhibitors (Fig 2A). Surprisingly, when we screened the same compound library for inhibition of PL^pro^ we identified only one compound that initially made the 10% cutoff, but this compound proved to be a false positive and we therefore ended up with no viable lead molecules for PL^pro^ (Fig 2B). An explanation for the absence of PL^pro^ lead scaffolds in our library most probably relates to the DUB like character of the protease together with its extremely narrow substrate specificity. The six validated M^pro^ hits can be categorized based on their electrophile class into aza-epoxyketones, chloro- and acyloxymethylketones and chloroacetamides (Fig 2C). We measured the kinetic inhibition parameter k_inact_/K_I_, for each compound (Fig 2D, Fig S4) and found that the aza-peptide epoxide, JCP474, was the most potent inhibitor of M^pro^ with a k_inact_/K_I_ value of 2526 ± 967 mol·sec^−1^. Interestingly, this compound was previously identified as a covalent inhibitor of SARS-CoV-1 M^pro^ (k_inact_/K_I_: 1900 ± 400 mol·sec^−1^)^30^.

**Figure 2.**
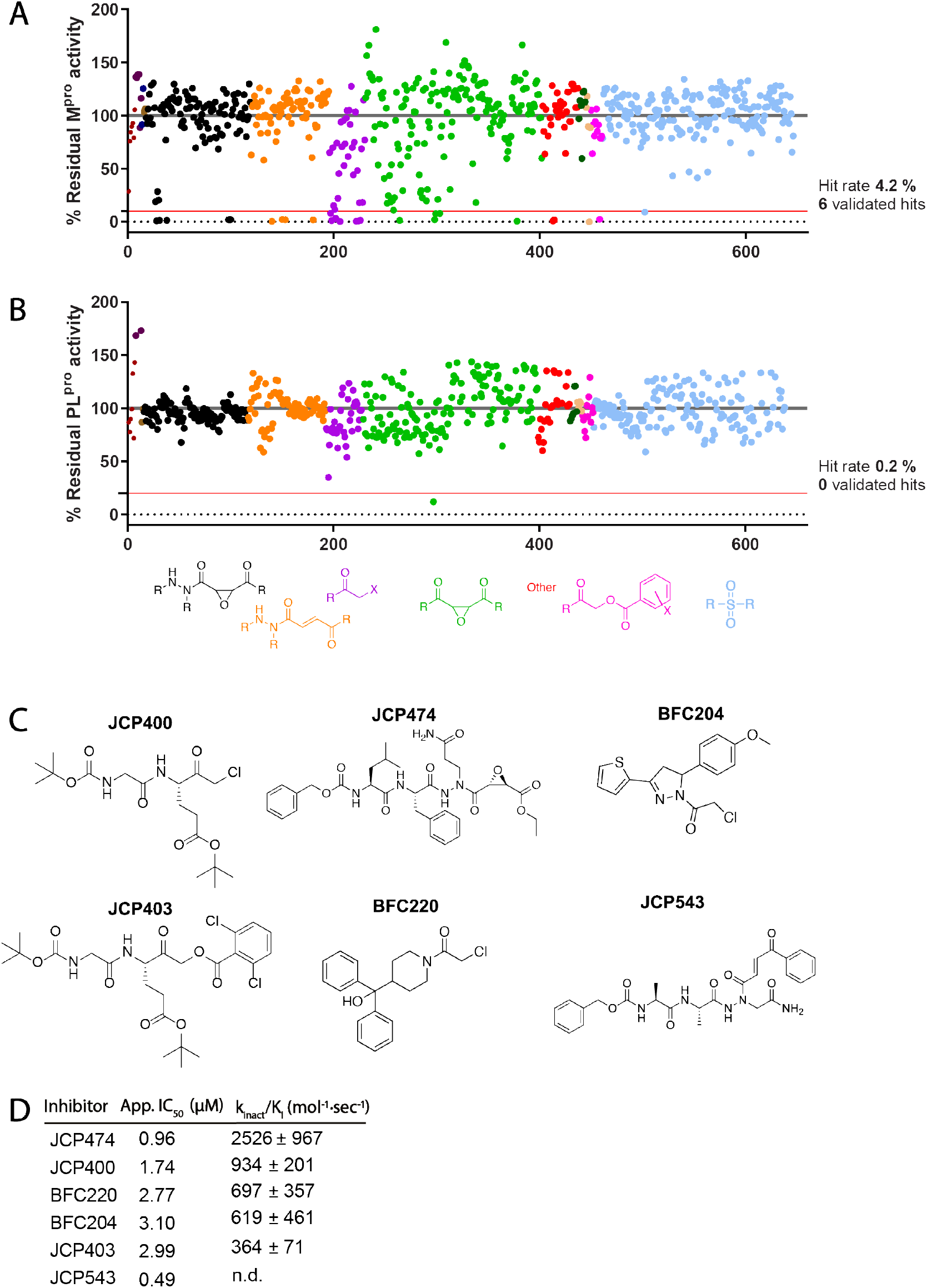
Screening of covalent inhibitor library against SARS-CoV-2 M^pro^ and PL^pro^. Residual activity of A) M^pro^ and B) PL^pro^ after 30 minutes incubation with 20 μM of each compound measured by cleavage rate of M^pro^ substrate 2 and Ac-LRGG-ACC for PL^pro^. C) Structures of M^pro^ hit compounds and D) their kinetic inhibition values measured without preincubation. Data are means ± SD of at least two replicate experiments

To probe the therapeutic potential of our M^pro^ inhibitors, we tested all of the compounds for inhibition of SARS-CoV-2 infection using a cellular model. A number of different types of host cells have been used in SARS-CoV-2 infection assays, with the most common cell type being Vero E6 cells of primate origin. However, as Vero E6 cells are not an accurate mimic of the human airway and lung epithelial cells that are the primary site of SARS-CoV-2 infection, we chose to instead use the lung adenocarcinoma cell line A549. This cell line is a more relevant lung derived human cell system, but it lacks sufficient expression of the ACE2 receptor to allow efficient infections by SARS-CoV-2^20^. Hence, we stably expressed the ACE2 entry receptor in A549 cells and achieved high level infection (typically greater than 50% infection) and replication during a 24-hours observation period. Using this cell system, we found that only two out of the six initial lead compounds blocked viral replication in these cells (Fig 3A). The two most potent inhibitors of M^pro^ *in vitro*, JCP474 and JCP543 were inactive in the cellular infection assay, likely due to the fact that they are both tripeptides with a polar P1 glutamine or asparagine residue resulting in poor cell permeability. The only two compounds that demonstrated activity were the chloromethylketone JCP400 and the acyloxymethylketone JCP403. These compounds showed relatively weak potency with greater than 75% inhibition only when applied at concentrations above 20 μM, which is well below cytotoxic concentrations (Fig. S5). This drop in potency of compounds in the cellular infection assay is consistent with what has been reported for other M^pro^ inhibitors^2,13^, and is likely due to poor cellular uptake and the difficulty in achieving complete inhibition of M^pro^ inside the host cell.

**Figure 3.**
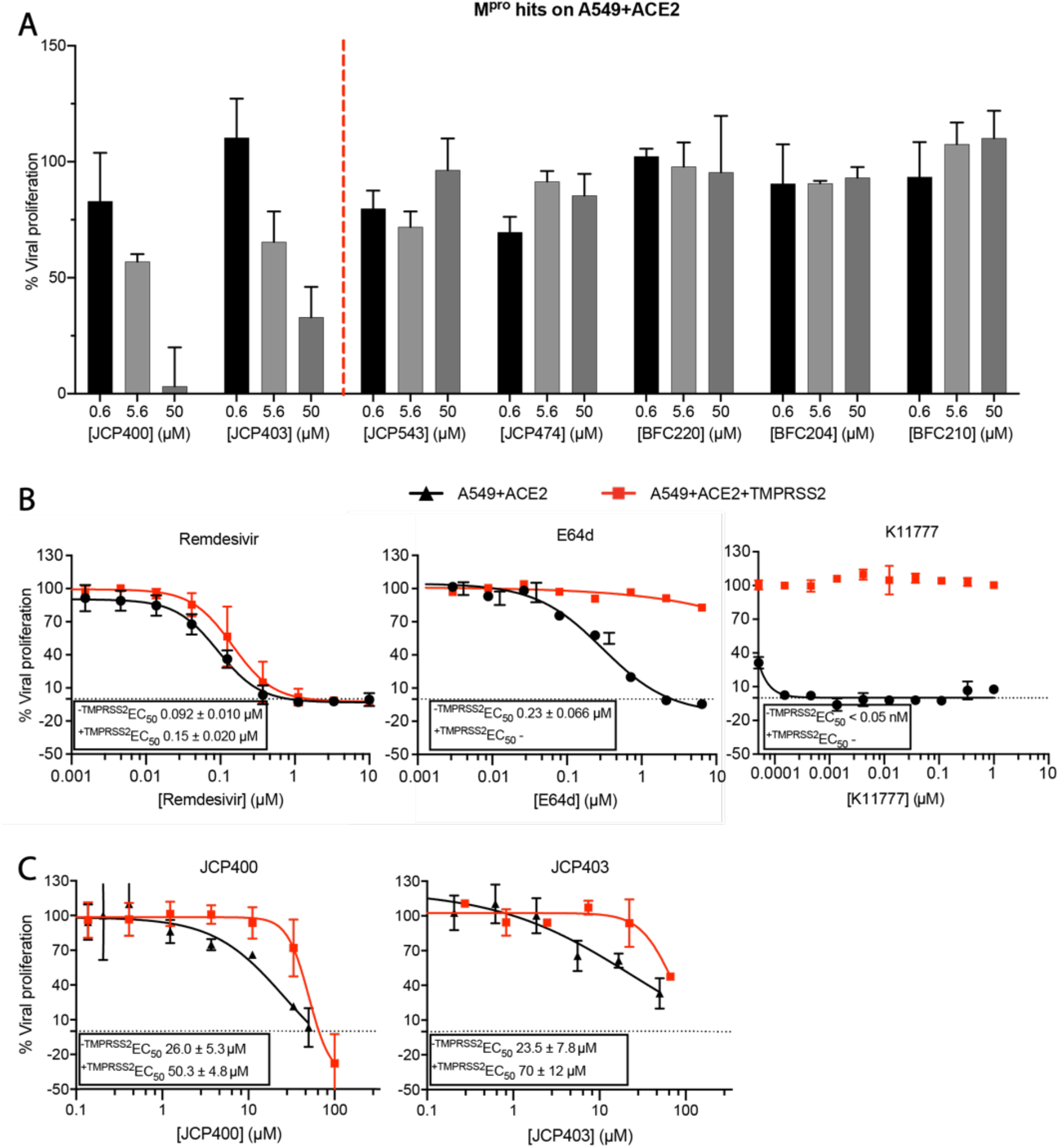
Potency of M^pro^ hits in cellular SARS-CoV-2 infection assays. A) Two out of six newly identified M^pro^ inhibitors are active in A549+ACE2 infection model. B) SARS-CoV-2 inhibition curves of Remdesivir, E64d and K11777 in A549+ACE2 cells with or without expression of TMPRSS2. C) SARS-CoV-2 inhibition curves of M^pro^ inhibitors JCP400 and JCP403 in A549+ACE2 cells with or without expression of TMPRSS2. Data are means ± SD of two replicate experiments.

One of our concerns about screening for M^pro^ inhibitors in our cysteine protease inhibitor library was the potential for hits to have cross-reactivity with other cysteine proteases. This becomes particularly problematic if compounds are only active against the virus at relatively high concentrations. The most likely family of off-target host proteases are the cysteine cathepsins, which are broadly expressed in many cell types and are accessible to small molecule and peptide-based inhibitors because of their lysosomal localization. Furthermore, recent studies have shown that SARS-CoV-2 can utilize multiple entry pathways into the host cell that depend on a variety of cellular proteases among which are cathepsin B and L, TMPRSS2 and furin^20,21^. One of the primary routes involves processing of the viral spike protein by the TMPRSS2 protease. This pathway is highly redundant with a pathway involving processing by cathepsin L (recent work has shown that Cat B is unable to independently process the spike protein^31^). Therefore, cathepsin inhibitors such as E64d and K11777 are highly potent inhibitors of viral entry in some cell lines but this activity is lost upon expression of TMPRSS2^20^. Hence, we sought to assess if either of our two lead M^pro^ inhibitors were active in the cellular assay as a result of inhibition of host cathepsins rather than as a result of inhibiting the virus encoded M^pro^ enzyme.

To address this issue, we generated A549+ACE2 cells that also express TMPRSS2, which is not expressed to a detectable level in regular A549 cells (data not shown), and investigated if expression of this alternate protease resulted in any change in antiviral activity. We first tested remdesivir and E64d and found that remdesivir was equipotent in both cell lines, while E64d completely lost its potency upon expression of TMPRSS2, consistent with previous studies^20^ (Fig 3B). Following a recent report showing that the cathepsin inhibitor K11777 is a highly potent SARS-CoV-2 antiviral compound^31^, we included this molecule in our analysis and found that it too lost all of its activity upon expression of TMPRSS2. For our two lead M^pro^ inhibitors, we found that their apparent EC_50_ values dropped by two to three-fold upon expression of TMPRSS2 (Fig 3C). Notably, both compounds displayed some signs of cytotoxicity at concentrations above 50 μM (Fig S5).

To confirm that the observed drop in potency of lead molecules upon TMPRSS2 expression was due to off-target reactivity of the compounds with cysteine cathepsins, we performed competition inhibition studies using the covalent cathepsin activity-based probe (ABP) BMV109. This ABP has been used to quantify levels of cathepsin activity in various cell-based systems^32–36^. Using this labeling approach, we found that JCP400 and JCP403 are both able to compete with BMV109 labeling of Cat B and L in A549+ACE2 cells (Fig 4A). As further validation of the off-target activity of the two lead molecules, we also tested the compounds for their ability to inhibit purified Cat B and L enzymes. These results confirmed that both are relatively potent inhibitors of cathepsins with IC_50_ values in the low micromolar range (Fig 4A-B).

**Figure 4.**
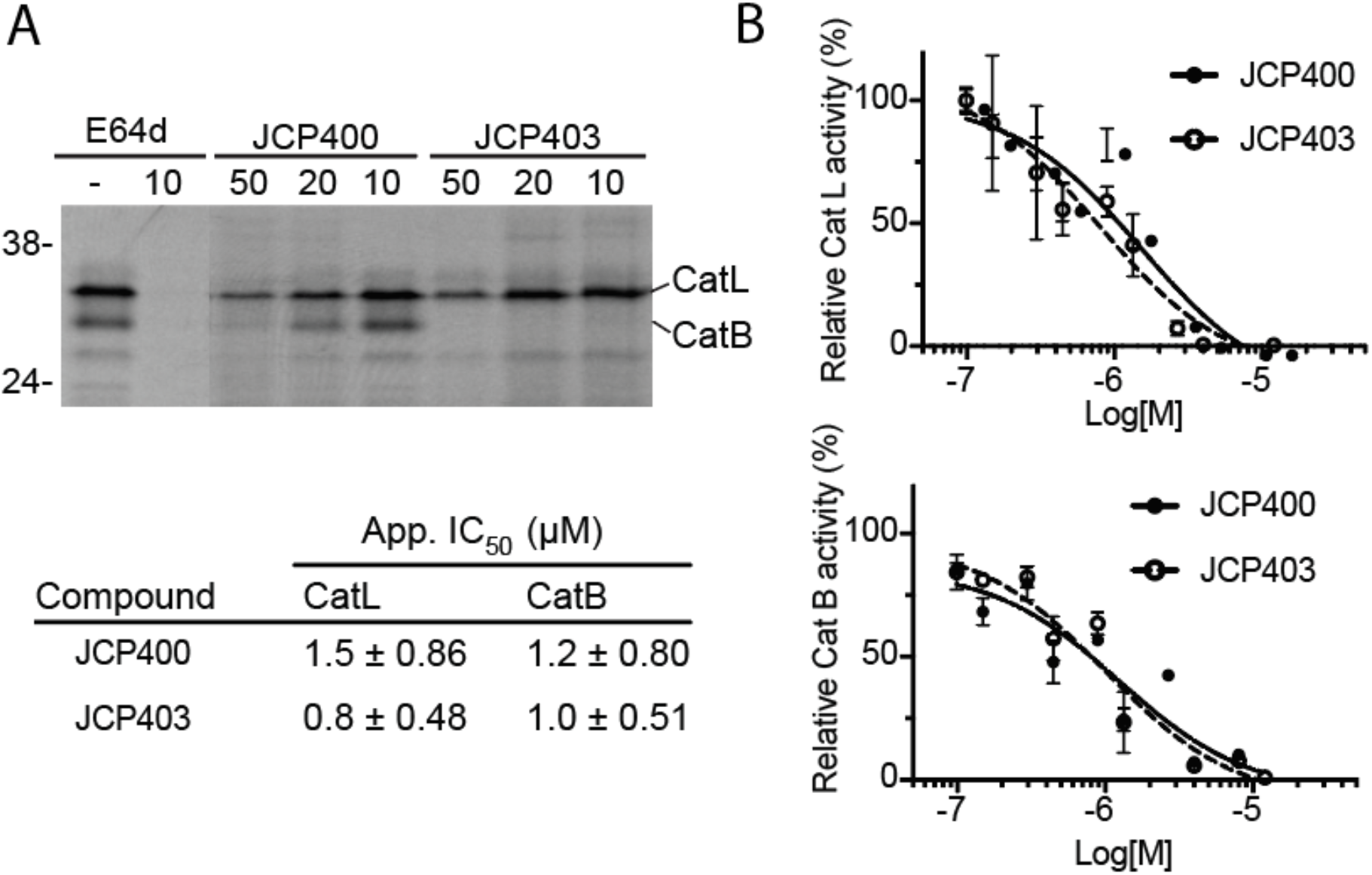
JCP400 and JCP403 inhibit cathepsin L and B. A) JCP400 and JCP403 compete with covalent labeling of broad spectrum cathepsin ABP BMV109 in A549+ACE2 cells. Cells were incubated with each compound for 1 h prior to addition of BMV109 B) JCP400 and JCP403 inhibit substrate cleavage of recombinant cathepsin L and B. Data are means ± SD of two replicate experiments.

Having confirmed that our newly identified compounds were cross-reactive with host cathepsins and that this activity was responsible for the bulk of their antiviral activity, we questioned whether previously reported M^pro^ inhibitors might have similar properties. We first evaluated five reported M^pro^ inhibitors for inhibition of human recombinant Cat B and L using a fixed time point fluorogenic substrate *in vitro* assay (Fig 5A). Surprisingly, the three aldehyde-containing inhibitors **GC373**, **11a**, and **11b** were highly potent with nanomolar IC_50_ values for both Cat L and Cat B. **Rupintrivir,** on the other hand, displayed no inhibition toward Cat B (tested up to 250 μM) and had only weak micromolar activity against Cat L.

**Figure 5.**
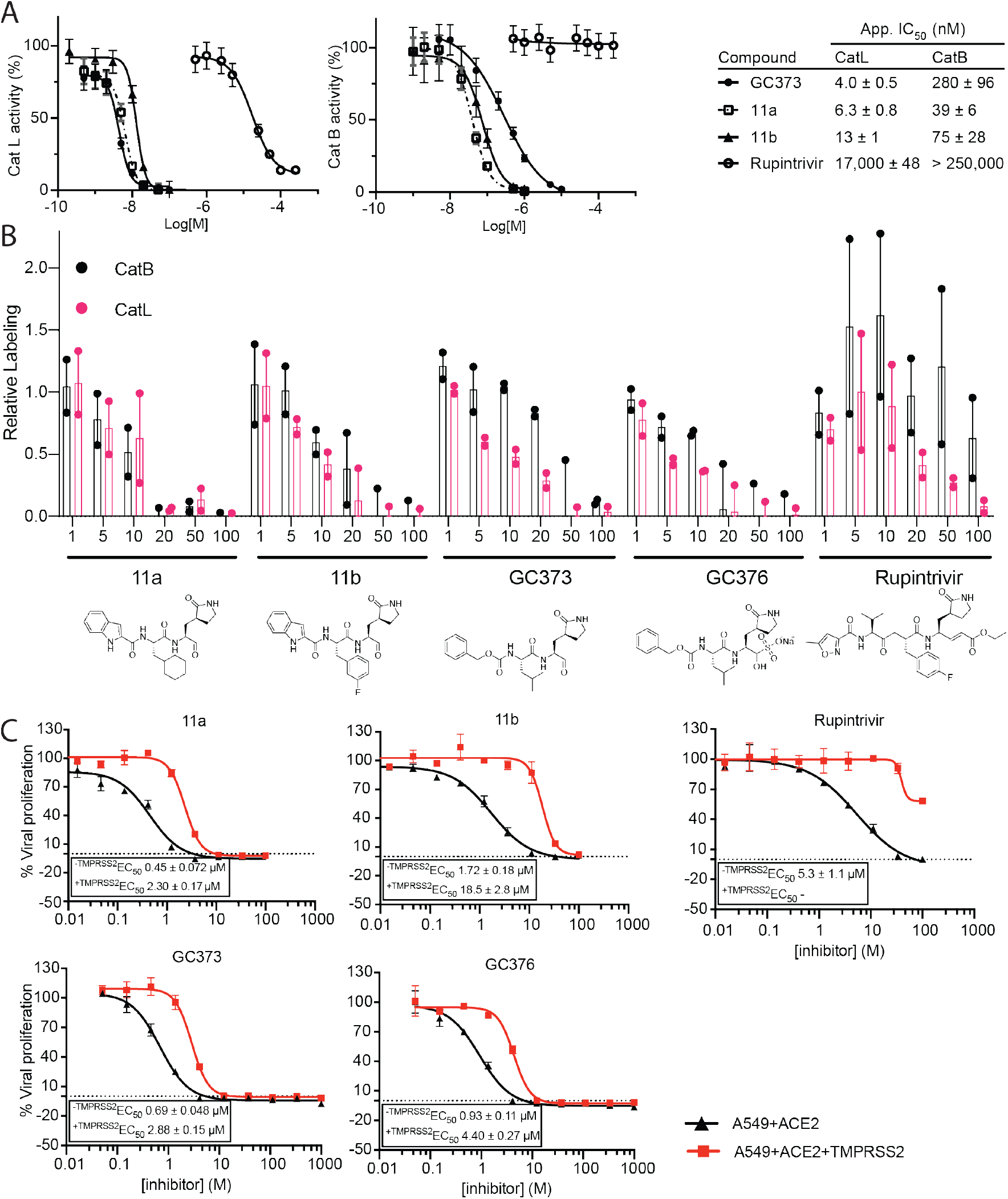
Reported M^pro^ inhibitors cross react with cathepsin B and L. A) Inhibition of recombinant cathepsins. Protease was incubated for 10 min with inhibitor prior to addition of substrate 6QC and fluorescent readout. Data are means ± SD of two replicate experiments. B) In-cell competition labeling with BMV109. A549+ACE2 cells were subjected to 1h treatment with inhibitor at indicated concentrations followed by 1h incubation with 1 μM BMV109. Cells were lysed and ran on SDS-PAGE gels that were scanned for in-gel fluorescence. Bar graphs represent relative densitometric quantification of two replicate experiments ± SD. C) Plots of EC_50_ curves of reported M^pro^ inhibitors in A549+ACE2 cells +/− TMPRSS2. Data are means ± SD of two replicate experiments.

We next evaluated whether the inhibitors were active against Cat B and L in A549+ACE2 cells. In-cell competition of the selected compounds with cathepsin labeling by BMV109 demonstrated that all of the reported M^pro^ inhibitors modified the active site residues of Cat B and L (Fig 5B, Fig S6). Consistent with the recombinant enzyme data, compounds **11a** and **11b** were active against cellular Cat B and L in the micromolar range with complete competition at 20 μM. The inhibitor **GC373** and its pro-drug form **GC376** show similar competition of Cat L between 5-10 μM and were slightly less potent toward Cat B with competition beginning between 20-50 μM. **Rupintrivir** was active against Cat L starting at 20 μM and showed only slight inhibition of Cat B labeling even at 100 μM.

Finally, we tested the reported M^pro^ inhibitors for activity in the A549+ACE2 cells with and without expression of TMPRSS2 to determine if cross reactivity with cathepsins was contributing to their antiviral activity. Indeed, we found that all five inhibitors showed a loss in potency upon TMPRSS2 expression similar to what we observed for our newly identified M^pro^ inhibitors. The effect appeared to be most prominent for aldehyde **11b**, which showed an 11-fold drop in potency. Interestingly, the α,β-unsaturated ketone **rupintrivir,** which has low micromolar activity in the cells lacking TMPRSS2, completely lost its antiviral activity when TMPRSS2 was expressed even though it showed minimal cathepsin cross reactivity (Fig 5B). Together with a lack of inhibitory activity against recombinant M^pro^ (Fig S7), this strongly suggests that **rupintrivir** derives all of its activity in cellular assays from weak inhibition of Cat L or possibly activity against other redundant proteases that can process the RBD to facilitate viral entry. The other compounds **11a**, **GC373** and **GC376** displayed a 4-5-fold decrease in potency upon expression of TMRPSS2 in the host cell (Fig 5C). Taken together, these results suggest that all of the tested M^pro^ inhibitors have some level of antiviral activity that is due to inhibition of host derived cathepsins and which is overcome to varying degree by the use of an alternate spike protein processing pathway employed by SARS-CoV-2.

In conclusion, inhibition of the M^pro^ and PL^pro^ proteases is considered to be a potentially viable therapeutic strategy for the treatment of COVID-19. However, because animal models of SARS-CoV-2 infection are still being optimized and controversy remains about cell systems that most accurately mimic aspects of the human infection (likely including viral entry pathways), it will be critical to assess key parameters of target selectivity of drug leads prior to clinical testing in humans. Furthermore, variability within the cellular systems used for antiviral testing can lead to flawed conclusions about lead candidate efficacy. The majority of current approaches only use inhibition of viral replication as a metric for efficacy of lead molecules without any direct confirmation of target inhibition. Only recently, has inhibition of processing of a genetically expressed M^pro^ substrate or labeling of active M^pro^ enzyme been established as a measure of M^pro^ activity in cells^26,37^. In this work, we describe our efforts to screen a library of approximately 650 diverse covalent inhibitor scaffolds against the two primary SARS-CoV-2 cysteine proteases, M^pro^ and PL^pro^. We failed to identify any inhibitors of PLpro and ultimately found only two inhibitors of M^pro^ that exerted antiviral activity in cell infection models, but only at relatively high concentrations. However, we found that the antiviral activity of these lead molecules as well as several previously reported M^pro^ inhibitors was related to their ability to inhibit host cathepsins, thus highlighting the importance of understanding compound selectivity and verifying target engagement. Taken together, our results point out the challenges for developing inhibitors of SARS-CoV-2 proteases and suggest that using strategically chosen cell lines for antiviral testing can help to prevent selection of compounds whose mechanisms of action can be easily overcome by redundant viral entry pathways. We strongly believe that our findings are of particular importance in light of drugs that are widely suggested for advancement into clinical trials such as **rupintrivir**^38,39^, or even have entered clinical trials such as K11777 (Selva Therapeutics, received FDA authorization for IND) and PF-07304814.

## METHODS

### M^pro^ expression and purification

Recombinant M^pro^ and the expression plasmid were gifts from D. Nomura (Berkeley). Expression and purification were performed as previously described for M^pro^ from SARS-CoV^40^ and SARS-CoV-2^13^. The gene encoding M^pro^ was synthesized and cloned into the pGEX vector resulting in a GST-M^pro^-6xHis fusion construct (pGEX-M^pro^), with the native M^pro^ cut site between GST and M^pro^ and a PreScission protease cut site between M^pro^ and the 6xHis tag. During expression, the N-terminal GST fusion is autoproteolytically cleaved by M^pro^ to yield the native N-terminus of the protease. Cleavage by 3C protease during purification yields the native C-terminus.

*E. coli* BL21 (DE3) were transformed with pGEX-M^pro^ and cultured in 1 L of 2xYT medium with ampicillin (100 μg/mL) at 37 °C. When the culture reached an OD_600_ of 0.8, protein expression was induced by addition of isopropyl-D-thiogalactoside (IPTG) to a final concentration of 0.5 mM. Expression was allowed to continue for 5 h at 37 °C. Cells were collected by centrifugation, resuspended in Buffer A (20 mM Tris, 150 mM NaCl, pH 7.8), and lysed by sonication. The lysate was clarified by centrifugation at 15,000 x g for 30 min at 4 °C. Clarified lysate was purified by NiNTA affinity using a HisTrap FF column (Cytiva). After loading the lysate, the column was washed with Buffer A and then protein was eluted over a gradient from Buffer A to Buffer B (20 mM Tris, 150 mM NaCl, 500 mM imidazole, pH 7.8). 3C protease was added to pooled elution fractions, and the mixture was dialyzed overnight at 4 °C into Buffer C (20 mM Tris, 150 mM NaCl, 1 mM DTT, pH 7.8). The dialyzed mixture was passed over a HisTrap FF column to remove the cleaved HisTag fragment and the His-tagged 3C protease. M^pro^ eluted in the flowthrough and was concentrated and buffer exchanged to Buffer D (20 mM Tris, 150 mM NaCl, 1 mM EDTA, 1 mM DTT, pH 7.8) using an Amicon 10kDa spin filter. Purified M^pro^ was aliquoted and stored at −80 °C.

### PL^pro^ expression and purification

For cloning of PL^pro^ with an N-terminal 6xHis-SUMO1 tag, synthetic fragments of the Nsp3 coding sequence derived from the original Wuhan strain were purchased from BioCat. and inserted into pUC57. The amino acid sequence of PL^pro^ (amino acids 1524-1883) was identified based on a homology blast using SARS-CoV-1 as template. The PL^pro^ sequence was amplified from pUC57-NSP3-BsaI-free-Fragments 1 and 2. The PCR products for SUMO1, PL^pro^ fragment 1 and PL^pro^ fragment 2, containing overlapping overhangs with unique restriction sites, were mixed in equimolar amounts and ligated into a linearized pet28a vector, resulting in a 6xHis-SUMO1-PLpro construct.

PCR primers:

fw SUMO1w/AgeImut: AATTCGAGCTCATGTCTGACCAGGAGGCA
rev SUMO1w/AgeImut: TCCTCACACCACCGGTTTGTTCCTGATAAACTTCAATCACATC
fw proPLfrag1: TCAGGAACAAACCGGTGGTGTGAGGACCATCAAGGTG
rev proPLfrag1: CAGAAAGCTAGGATCCGTGGTGTGGTAGT
fw proPLfrag2: ACCACACCACGGATCCTAGCTTTCTGGGCAGG
rev proPLfrag2: GTGCGGCCGCAAGCTTTCACTTGTAGGTCACAGGCTTGA

*E. coli* BL21 (DE3) were transformed with 6xHis-SUMO1-PL^pro^ in pet28a. Cells were grown in 2 liters LB medium supplemented with 50 μg/mL kanamycin at 37 °C. At an OD_600_ of 0.6, cells were further diluted with precooled LB medium supplemented with kanamycin, to a final volume of 4 liters. Protein expression was induced by addition of 0.1 mM IPTG and 0.1 mM ZnSO_4_ at 18 °C. Following a 24 h induction, cells were harvested by centrifugation and resuspended in Lysis buffer (50 mM NaH_2_PO_4_, 300 mM NaCl, 10 mM Imidazole pH 8.0, 1 mM ß-Mercaptoethanol), followed by sonication. Cell lysates were incubated with 50 μg / ml DNase I and 1 mM MgCl_2_, at 4 °C for 45 min, and subsequently subjected to ultracentrifugation at 100,000 x g for 1 h at 4 °C. Next, the clarified lysates were incubated with NiNTA beads (Qiagen) to capture 6xHis-SUMO1-PL^pro^. Following extensive washing, GST-SENP was added to the beads to cleave at the C-terminus of SUMO1, resulting in elution of untagged PL^pro^ with the native N-terminus. GST-SENP was captured and removed from the eluate by incubation with GSH beads (GE Healthcare). Purified PL^pro^ was further concentrated using an Amicon 5 kDa spin filter to a final concentration of 1 mg/ml and stored at −80 °C (50 mM NaH_2_PO_4_ pH 8.0, 300 mM NaCl). Proteolytic activity of purified PL^pro^ against Z-LRGG-AMC was tested over time. Prior to activity assays, PL^pro^ was activated by incubation in reaction buffer (150 mM NaCl, 20 mM Tris-HCl pH 7.5, 0.05% Tween-20, 0.2 mg/mL Ovalbumin) containing 5 mM DTT. RFU values were measured immediately in an Enspire Plate Reader (PerkinElmer). Each dot represents the mean of 3 independent experiments.

### Virus stock

The SARS-CoV-2 isolate used in this study was derived from a patient at Heidelberg University Hospital. This Heidelberg strain was passaged in VeroE6 cells, aliquoted and stored at −80° C. Virus titer was measured by plaque assay in VeroE6 cells.

### Cell lines

VeroE6 and A549 cells were obtained from American Type Culture Collection (ATCC) and tested at regular intervals for mycoplasma contamination. Generation and cultivation of A549 cells stably expressing ACE2 (A549+ACE2) was described recently^41^. A549+ACE2 cells stably expressing the TMPRSS2 protease were generated by lentiviral transduction. Lentivirus stocks were produced by transfection of HEK293T cells with a pWPI plasmid encoding for TMPRSS2 and the pCMV-Gag-Pol and pMD2-VSV-G packaging plasmids (kind gifts from D. Trono, Geneva). Two days after transfection, supernatant containing lentiviruses were collected, filtered through a 0.44 μm pore size filter, and used for transduction of A549+ACE2 cells followed by selection with 2 μg/mL puromycin. For all viral infection assays, the cells were cultured in Dulbecco’s modified Eagle medium (DMEM, Life Technologies) containing 10% or 20% fetal bovine serum, 100 U/mL penicillin, 100 μg/mL streptomycin and 1% non-essential amino acids (complete medium). For all other assays, A549+ACE2 cells were cultured in Roswell Park Memorial Institute (RPMI, Corning, REF: 10-040-CV) 1640 medium containing 2 g/L of glucose, 0.3 g/mL of L-glutamine, and supplemented with 10 v/v% FBS, 100 units/mL penicillin, 100 μg/mL streptomycin, and 625 μg/mL of Geneticin (G418). All cells were grown in a 5% CO_2_ humidified incubator at 37 °C.

### Primary library screening

A compound library of 646 diverse molecules containing electrophilic warheads were kept in 1, 10 or 50 mM DMSO stock solutions at −80 °C for long-term storage. All assays were conducted in black, opaque flat squared bottom 384-well plates (Greiner Bio-One, REF: 781076) containing a final reaction volume of 50 μL. Assays volumes and concentrations used were as follows: 0.5 μL of 1 mM compound was added to the wells, followed by 25 μL of 200 nM M^pro^ or 150 nM PL^pro^ (M^pro^ buffer: 150 mM NaCl, 20 mM Tris pH 7.5, 1 mM EDTA, PL^pro^ buffer: 150 mM NaCl, 20 mM Tris pH 7.5, 0.05% Tween-20). After 30 min incubation at 37 °C, 24.5 μL of 20 μM M^pro^ substrate 2 or 100 μM Ac-LRGG-ACC (for PL^pro^) was added to the wells and the fluorescent measurement was started immediately. The final concentrations of compound, enzyme and substrate were 10 μM, 100 nM / 75 nM, 10 μM / 50 μM (M^pro^ / PL^pro^) respectively. Each 384-well plate contained at least 20 positive controls in which the compound was 10 μM ebselen for M^pro^ assays and heat inactivated (10 min, 95 °C) enzyme for PL^pro^ assays. Similarly, at least 20 negative controls were incorporated in each 384-well plate where 0.5 μL compound was swapped with 0.5 μL DMSO. Raw slope values were calculated as the slope of the absolute RFU versus time for the first 15 min of the experiment. Then, percentage activity was calculated by normalizing between slope values of the positive and negative controls. The inhibition threshold for M^pro^ was 90 % whereas for PL^pro^ a threshold of 80 % was chosen because of the low hit rate. All fluorescent measurements for substrates containing a sulfo-Cy5 or ACC moiety were read above the well with a Biotek Cytation3 Imaging Reader (7.00 mm read height, gain = 100, Cy5 = λ_ex_ 650 nm; λ_em_ 670 nm or ACC = λ_ex_ 355 nm; λ_em_ 460 nm, gain = 65, and normal read speed).

### IC_50_ value and kinetic parameter determination

Dose-response studies were performed by mixing 200 nM M^pro^ with 20 μM KS011 in a 384-well plate. Immediately before starting the fluorescence measurement, a dilution series of 6-10 different concentrations of inhibitor were added to the wells and the fluorescence intensity was recorded for 1 h. Apparent IC_50_ values were estimated by fitting the normalized linear slopes to eq 1 using a four-parameter fit. Using the same data, k_obs_ at each inhibitor concentration was estimated by nonlinear fitting of each progress curve to eq 2. The k_inact_/K_i_ could be determined by nonlinear fitting of eq 3 to k_obs_ as function of inhibitor concentration.

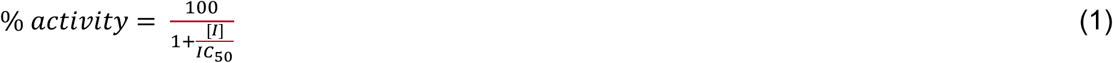

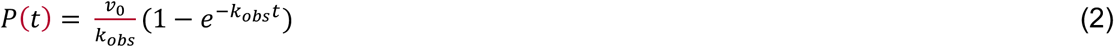

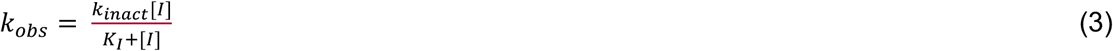

Inhibition studies with recombinant cathepsin B and L were performed with by incubating the serially diluted compounds for 10 min with either 40 nM Cat B or 10 nM Cat L (kind gifts from B. Turk, Ljubljana) in 50 mM citrate buffer (pH = 5.5, 5 mM DTT, 0.1% triton X, 0.5% CHAPS) and subsequent addition of 10 μM quenched-fluorescent substrate 6QC^42^. Fluorescence intensity was recorded using a plate reader at λ_ex_ 650 nm, λ_em_ 670 nm. Apparent IC_50_ values were determined similarly as for M^pro^ assays. All experiments were performed in duplicate. All data were analyzed using GraphPad Prism (v8.4).

### Competition assay in living A549 cells

In a 24-well plate, 1 μL of 200x inhibitor concentration was added to approximately 10^5^ A549+ACE2 cells in 200 μL medium containing 1% DMSO and incubated for 1 h at 37 °C. 1 μL of BMV109 was added at a final concentration of 1 μM and incubated for 1h. Medium was removed, and cells were detached from culture plate by incubating with 100 μL 0.05 % Trypsin, 0.5 mM EDTA solution for 10 min at 37 °C. Cells were spun down, washed twice with PBS and lysed by four succeeding freeze-thaw cycles via submersion of Eppendorf tubes in a 37 °C water bath and liquid nitrogen respectively. Protein concentration of lysate was determined using BCA assay, Laemmli’s sample buffer was added at 4-fold dilution and samples were boiled for 5 min before running them on 15% SDS-PAGE gel. In-gel detection of fluorescently labeled proteins was performed directly in the wet gel slabs on the Typhoon Variable Mode Imager (Amersham Biosciences) using Cy5 settings (λ_ex_ 650 nm, λ_em_ 670 nm). Densitometric analysis of protein bands on gels was performed using ImageJ (v1.52p).

### Antiviral assays

A549+ACE2+/−TMPRSS2 were seeded at a density of 1.5 × 10^4^ cells per well of a flat bottom 96-well plate (Corning). On the next day, for each compound serial dilutions of at least ten concentrations were prepared in complete DMEM and added to the cells. After 30 minutes, SARS-CoV-2 (MOI=1) is added into the compound containing medium. 24 h post infection, plates were fixed with 6% of formaldehyde and cells were permeabilized using 0.2% Triton-X100 in PBS for 15 min at room temperature. After washing with PBS and blocking with 2% milk in PBS / 0.02% Tween-20 for 1 hour at room temperature, cells were incubated with a double strand RNA-specific antibody (Scicons, Hungary) for 1 hour at room temperature. After three times washing with PBS, bound primary antibody was detected with a secondary antibody (anti-mouse IgG), conjugated to horseradish peroxidase. Bound secondary antibody was quantified using TMB (3,3’,5,5’-Tetramethylbenzidine) substrate (Thermo Fisher Scientific) and photometry at 450 nm in a plate reader. Background absorbance was measured at 620 nm. To determine cytotoxicity of the compounds, non-infected A549-derived cells were treated in the same way as the infected cells. After 24 hours, intracellular ATP content was quantified by using the CellTiter Glo^®^ Luminescent Cell Viability Assay (Promega) according to the instructions of the manufacturer. Values were normalized using solvent control (0.5% DMSO). Each experiment was performed in duplicate, and two independent biological replicates were conducted.

### Chemistry methods

All reactions were performed exposed to atmospheric air unless noted otherwise and with solvents not previously dried over molecular sieves or other drying agents. Reactions containing light sensitive materials were protected from light. The ACS reagent grade N,N’-dimethylformamide (DMF), molecular biology grade dimethyl sulfoxide (DMSO), and all other commercially available chemicals were used without further purification. Reaction temperatures above 23 °C refer to incubator temperatures, controlled by a temperature modulator. Reaction progress and purity analysis was monitored using an analytical LC-MS. The LC-MS systems used was either a Thermo Fisher Finnigan Surveyor Plus equipped with an Agilent Zorbax 300SB-C18 column (3.5 μm, 3.0 × 150 mm) coupled to a Finnigan LTQ mass spectrometer or an Agilent 1100 Series HPLC equipped with a Luna 4251-E0 C18 column (3 μm, 4.6 x 150 mm) coupled to a PE SCIEX API 150EX mass spectrometer (wavelengths monitored = 220, 254 and 646 nm). Purification of intermediates and final compounds was carried out using either a semi-preparative Luna C18 column (5 μm, 10 x 250 mm) attached to an Agilent 1260 Infinity HPLC system or a CombiFlash Companion/TS (Teledyne Isco) with a 4 or 12 g reverse phase C18 RediSep Rf Gold column (wavelengths monitored = 220 & 254 nm). Information regarding gradient programs for purifications can be found in the Chemistry Protocols section below. Intermediates were identified by their expected m/z using LC-MS. Rupintrivir was purchased from Tocris Bio-Techne. E64d was a gift from American Life Sciences Pharmaceuticals (to Christoph Peters) and Remdesivir was purchased.

### Synthesis of internally quenched and fluorogenic substrates

Fmoc-ACC-OH was synthesized as described^43^. Standard Fmoc chemistry was performed on Rink AM resin as described^42^. Internally quenched peptide substrate sequences were synthesized on 2-Chlorotrityl resin using standard Fmoc chemistry as previously described^44^. Peptides were cleaved from resin using 1,1,1,2,2,2-hexafluoroisopropanol to maintain the protecting groups on the amino acid side chains. After cleavage from the resin, Sulfo-Cy5-COOH (2 eq) was coupled to the free N-terminus by mixing with coupling reagent O-(1H-6-Chlorobenzotriazole-1-yl)-1,1,3,3-tetramethyluronium hexafluorophosphate (HCTU) (1.2 eq) and 2,4,6-collidene (1.2 eq) in DMF. The solution of activated acid was added to the amine and agitated at RT overnight. After the reaction went to completion according to LC-MS, the intermediate was purified using preparative reverse phase HPLC. After the purified product was collected and concentrated *in vacuo*, removal of protecting groups was achieved by dissolving the intermediate in 80:20 TFA:DCM and stirring at RT for 1 h. The reaction was then concentrated *in vacuo* and the crude material was used without further purification. Coupling of sulfo-QS21-Osu was achieved by dissolving the intermediate in DMSO and DIPEA (1.5 eq) and agitating for 24h at 37 °C. The reaction was then purified using preparative HPLC and fractions were collected and concentrated *in vacuo*. The residue was dissolved in 1:1 MeCN:H_2_O (0.1% TFA) and lyophilized to yield the M^pro^ substrate (1-4) as a blue powder.

## Data Availability

All data and information necessary to reproduce the results reported in this manuscript are provided. Any additional data that support the findings of this study is available upon reasonable request.

## ACKNOWLEDGEMENTS

We thank members of the Nomura lab at University of California, Berkeley for providing original stocks of the M^pro^ protease and plasmids; James Powers (University of Georgia), William Roush (The Scripps Research Institute) and Benjamin Cravatt (The Scripps Research Institute) for the directed irreversible inhibitors of cysteine proteases; Joanna Lemieux at University of Alberta for providing GC376 and GC373 compounds; Hong Liu at Shanghai Institute of Materia Medica for providing compounds 11a and 11b; Ania Plaszczyca for initial help to establish the in-cell ELISA SARS-CoV-2 detection protocol; members of the Boris Turk Lab at Jozef Stefan Institute for providing the recombinant cathepsin proteases used in this study.

## AUTHOR CONTRIBUTIONS

**Kas Steuten:** Conceptualization, Investigation, Formal Analysis, Visualization, Writing – original draft; **Heeyoung Kim:** Investigation, Resources; **John C. Widen:** Conceptualization, Investigation, Formal Analysis, Visualization, Writing – review & editing; **Brett M. Babin:** Conceptualization, Investigation, Formal Analysis, Writing – review & editing **Ouma Onguka:** Investigation, Resources**, Scott Lovell:** Conceptualization, Investigation; **Oguz Bolgi:** Resources, Investigation **Berati Certikan:** Investigation, Resources; **Mirko Cortese:** Resources; **Ryan Muir:** Resources; **John Bennett:** Resources; **Ruth Geiss-Friedlander:** Resources, Conceptualization, Supervision; **Christoph Peters:** Conceptualization, Funding Acquisition**, Ralf Bartenschlager:** Conceptualization, Supervision, Resources, Funding Acquisition, Writing – review & editing; **Matthew Bogyo:** Conceptualization, Funding Acquisition, Supervision, Writing – review & editing

