## Supplemental Figures for "Challenges for targeting SARS-CoV-2 proteases as a therapeutic strategy for COVID-19"

**Supporting Figures**

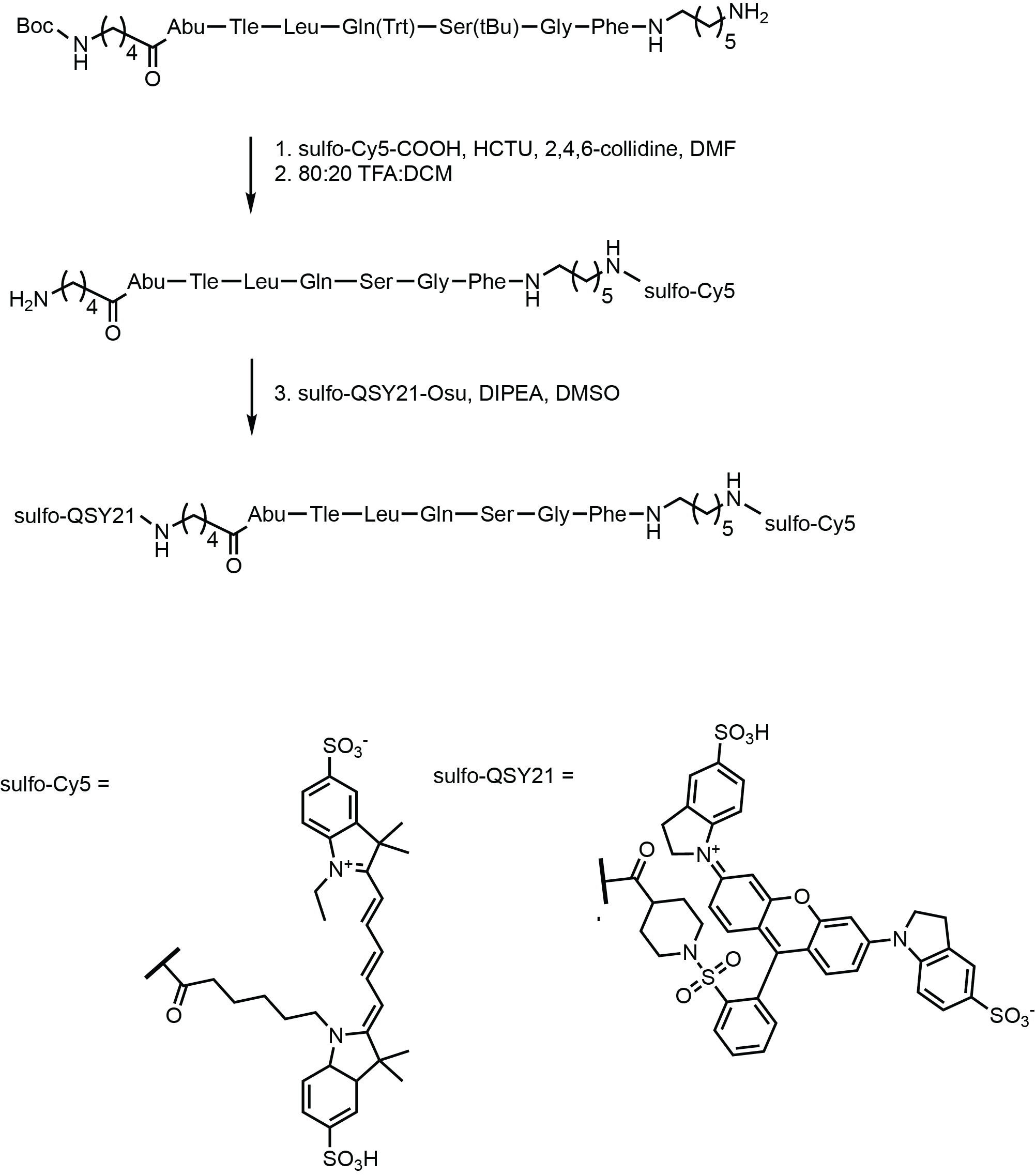

**Fig S1 Synthetic route to internally quenched fluorescent M^pro^ substrate 1.** Synthesis of other M^pro^ substrates followed a similar procedure. See methods section for synthetic details.

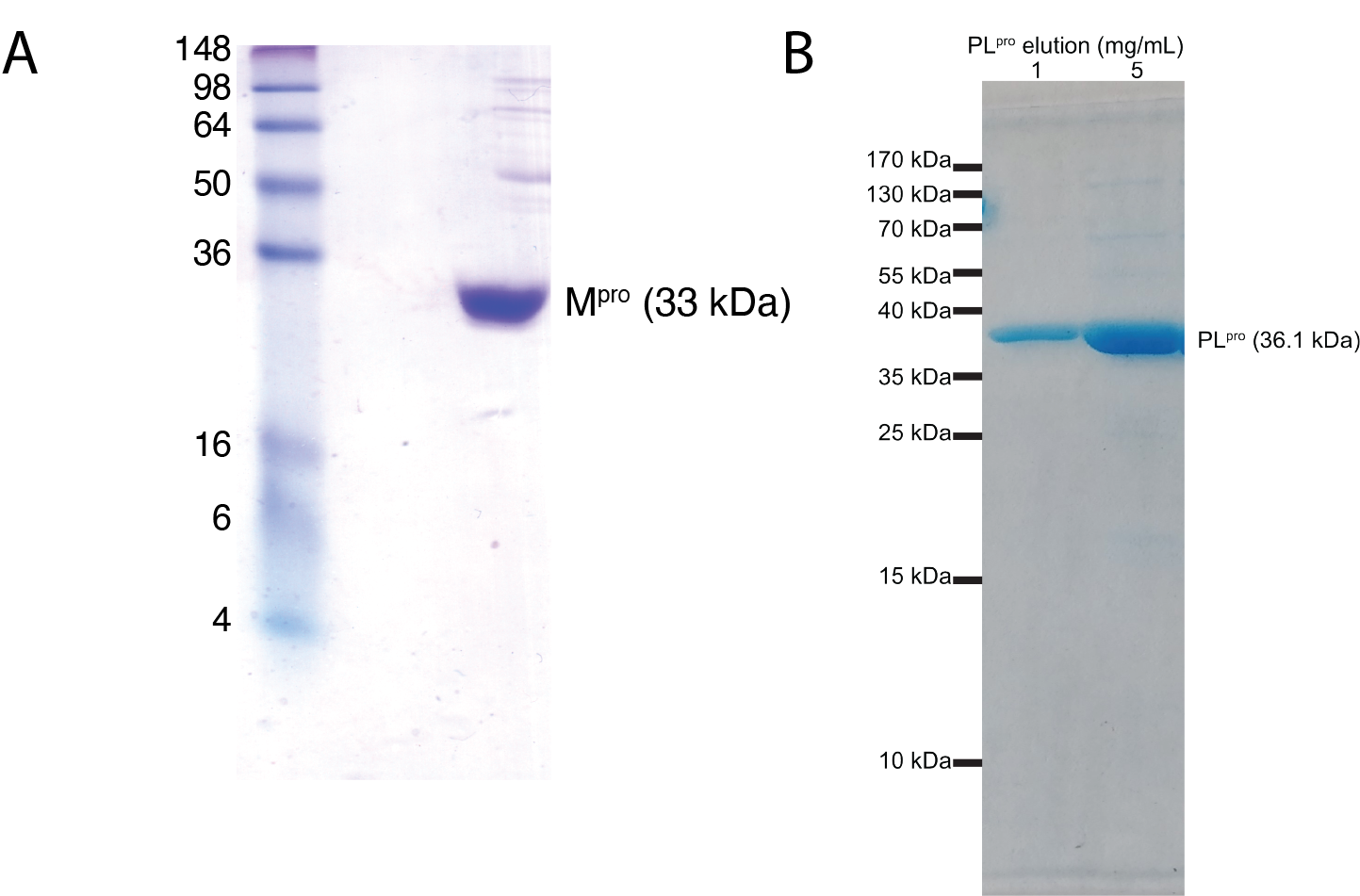
 **Fig S2 Purification gels of SARS-CoV-2 M^pro^ and PL^pro^.** A) Recombinant M^pro^ produced using a GST fusion construct. B) Recombinant PL^pro^ produced using a SUMO1 fusion construct. Images represent SDS-PAGE gels of purified recombinant proteases.

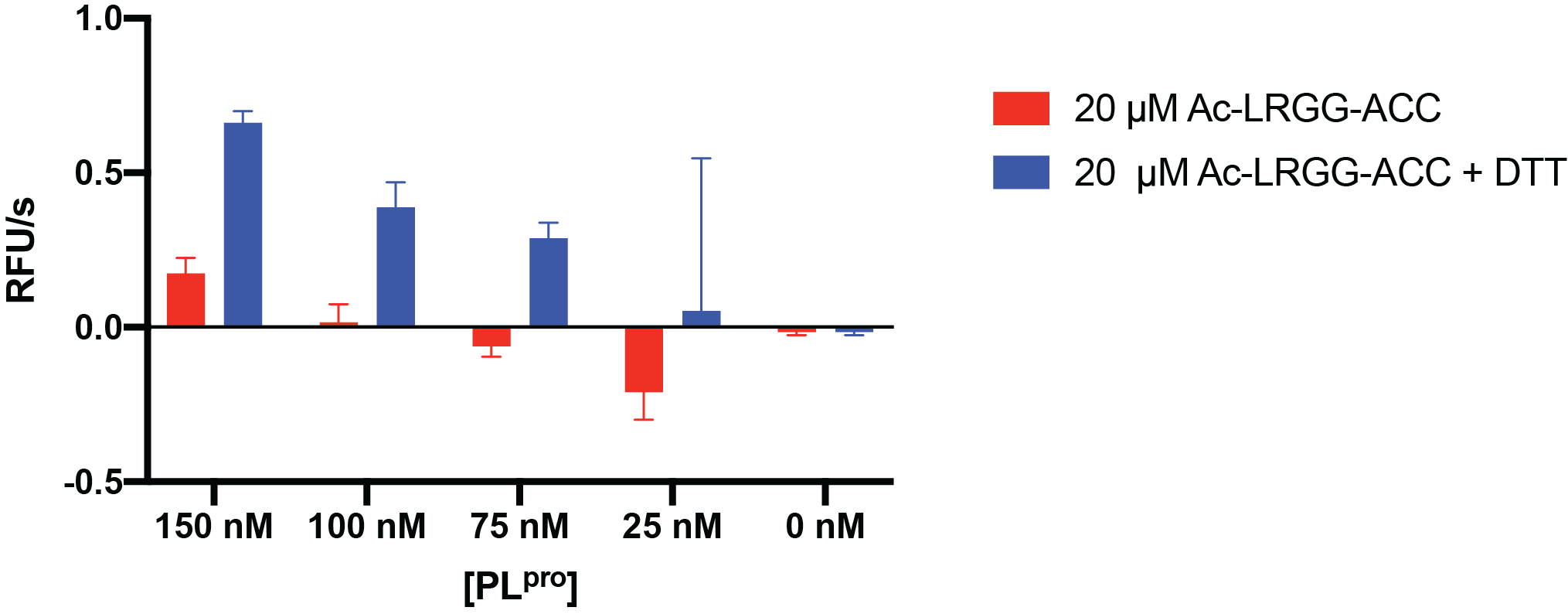

**Fig S3 2 mM DTT in buffer increases cleavage rate of Ac-LRGG-ACC by PL^pro^.** Cleavage rate was determined as slope of linear part of the fluorescence intensity progress curve. Data are means ± SD of three replicate experiments.

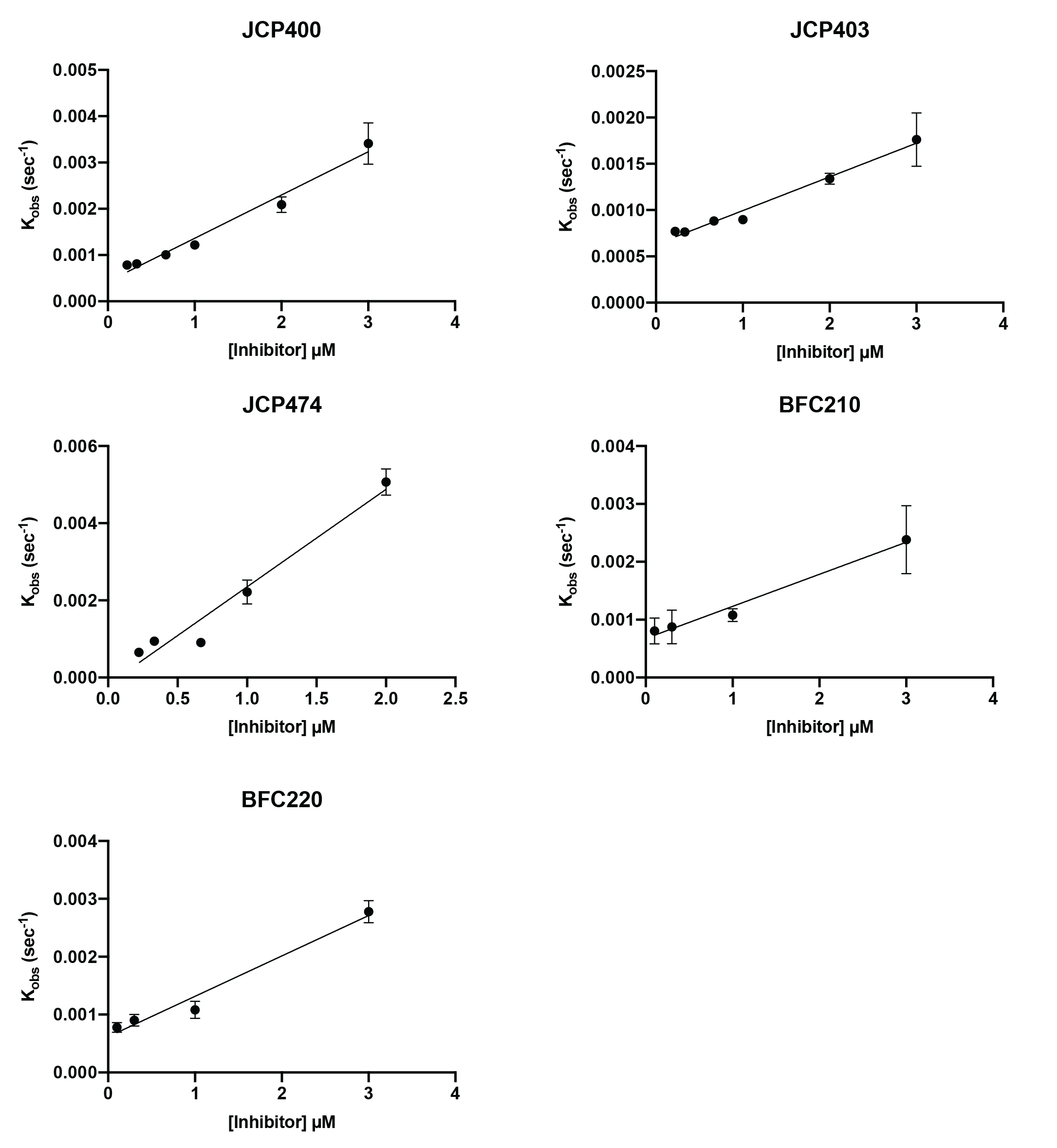

**Fig S4 Slopes used for determination of kinetic inhibition parameter k_inact_/K_I_ for six covalent M^pro^ inhibitors.** 10 µM of M^pro^ substrate 2 was mixed with indicated amounts of inhibitor and immediately after addition of 100 nM M^pro^ enzyme, the fluorescent intensity measurement was started using a plate reader. Straight lines are standard linear fits of which the slope corresponds to k_inact_/K_I_. Data are means ± SD of three replicate experiments.

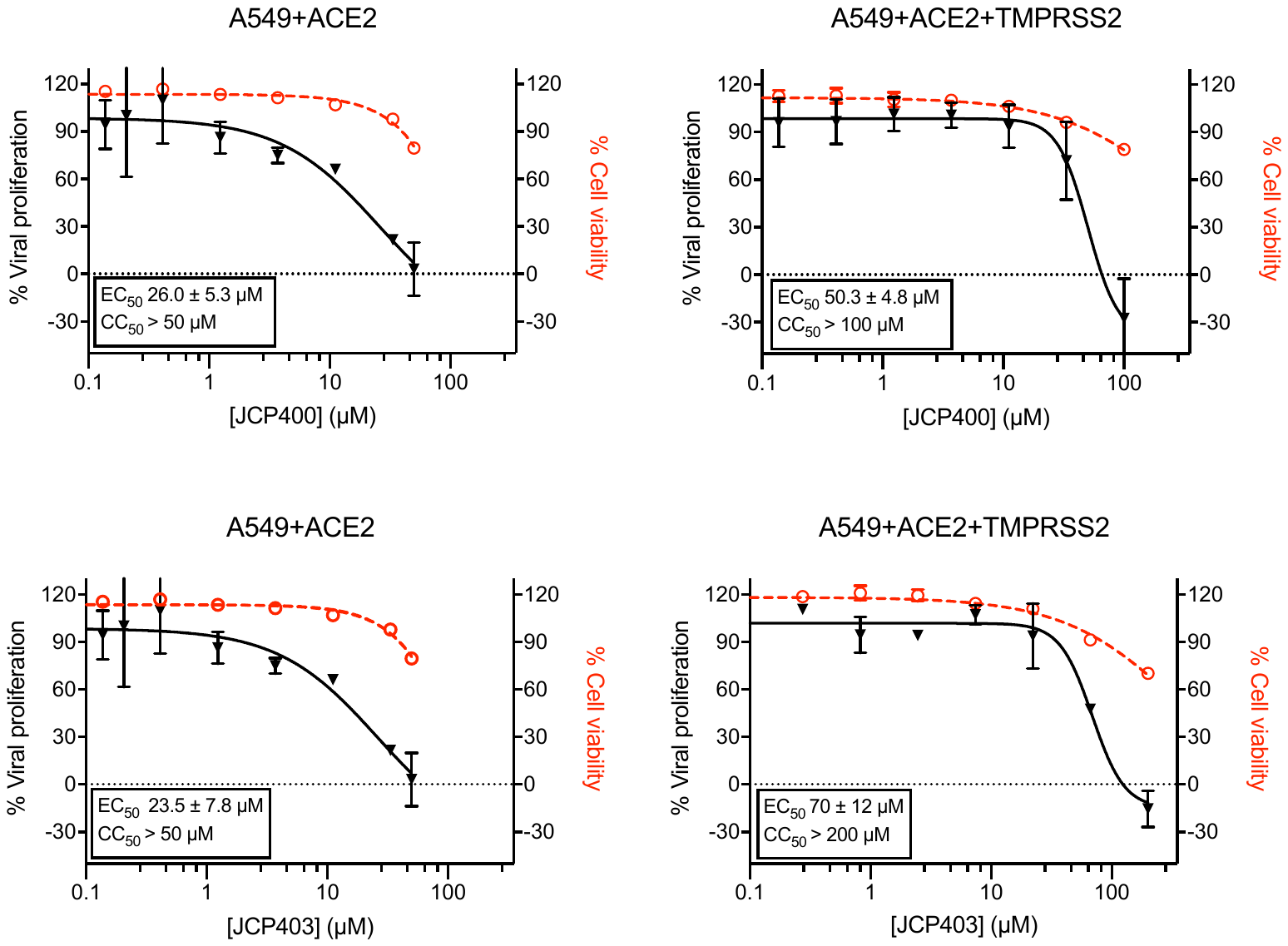

**Fig S5 Cytotoxicity profile of JCP400 and JCP403 in A549+ACE2+/-TMPRSS2 corresponding to main Figure 3B.** Cytotoxicity was measured using CellTiter Glo® Luminescent Cell Viability Assay as described in methods section. Data are means ± SD of two replicate experiments.

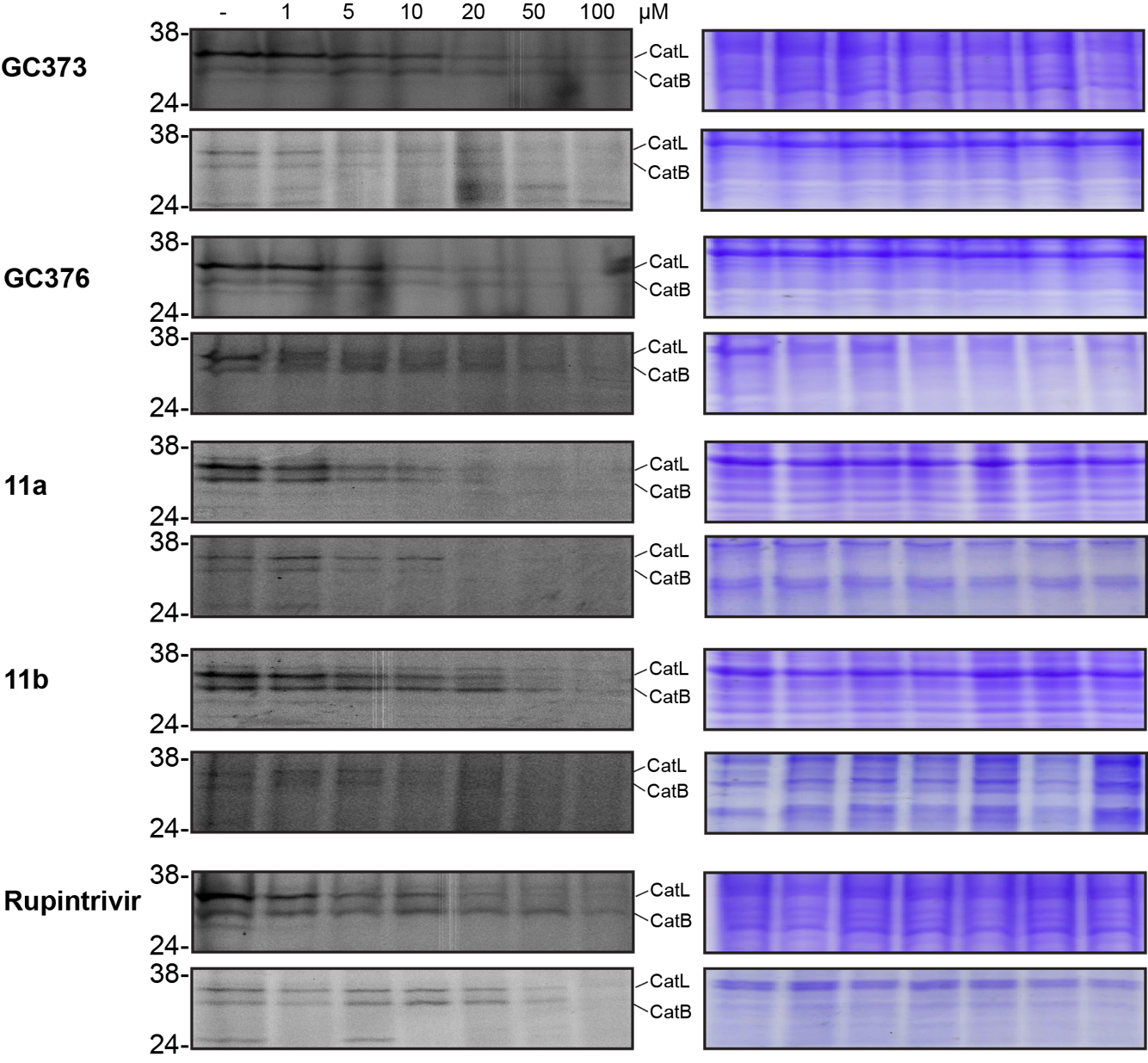

**Fig S6 Gel images of BMV109 competition experiments.** Shown are SDS-PAGE gels of A549+TMPRSS2 lysate that was incubated with indicated concentration of inhibitor for 1 h followed by 1 h incubation with BMV109 prior to washing and further sample preparation. In the right column, gel images after Coomassie staining are depicted to indicate equal protein loading.

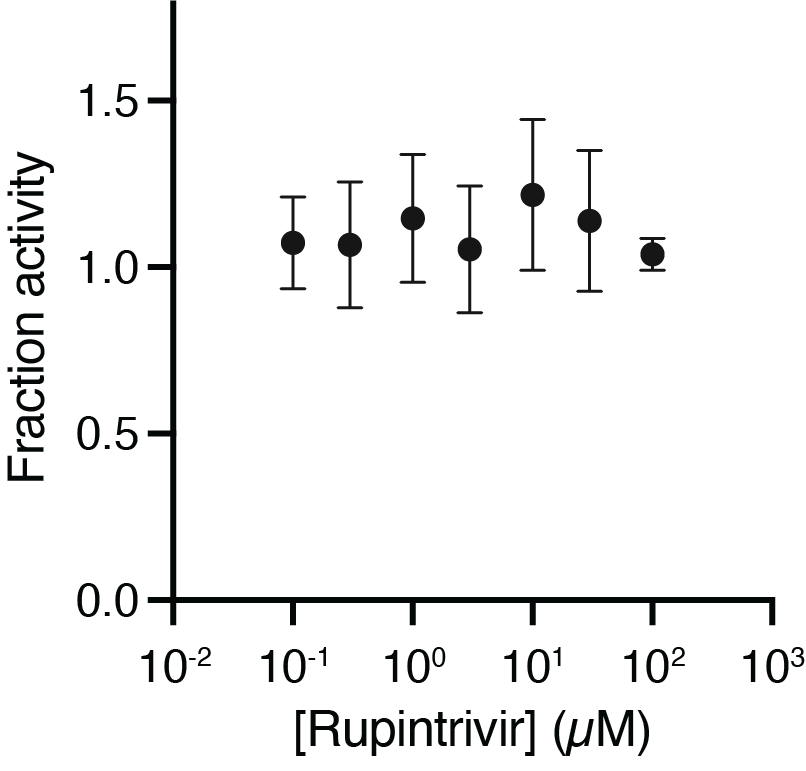

**Fig S7 Rupintrivir does not affect M^pro^ activity at concentrations up to 100 µM.** Inhibition was measured without preincubation using M^pro^ substrate 1.
